# First molecular phylogeny of the freshwater planarian genus *Girardia* (Platyelminthes, Tricladida) unveils hidden taxonomic diversity and initiates resolution of its historical biogeography

**DOI:** 10.1101/2022.07.19.500599

**Authors:** Lisandra Benítez-Álvarez, Ronald Sluys, Ana María Leal Zanchet, Laia Leria, Marta Riutort

**Affiliations:** Departament de Genètica, Microbiologia i Estadística and Institut de Recerca de la Biodiversitat (IRBio), Universitat de Barcelona, Avinguda Diagonal 643, 08028, Barcelona, Catalonia, Spain; Naturalis Biodiversity Center, P. O. Box 9517, 2300 RA Leiden, The Netherlands; Instituto de Pesquisas de Planárias and Programa de Pós-Graduacão em Biologia, Universidade do Vale do Rio dos Sinos (UNISINOS), 93022-750 São Leopoldo, Rio Grande do Sul, Brazil

**Keywords:** *Girardia*, evolutionary relationships, historical biogeography, hypogean diversity, introduced species, taxonomy, Tricladida, new species

## Abstract

The genus *Girardia* (Platyhelminthes: Tricladida) comprises several species of which some have spread from their original areas of distribution in the Americas to other parts of the globe. Due to great anatomical similarities between species, morphology-based phylogenetic analyses struggled to resolve the affinities between species and species-groups. This problem is exacerbated by the fact that populations of *Girardia* may show only asexual reproduction by fissiparity and, thus, do not exhibit a copulatory apparatus, which hampers taxonomic identification and extraction of phylogenetic characters. In the present work this problem has been resolved by constructing a molecular phylogeny of the genus. Although our samples do not include representatives of all known species, they cover a large part of the original distributional range of the genus *Girardia*. Our phylogenetic results suggest the presence of two main clades, which are genetically and karyologically highly differentiated. North and South American nominal *G. tigrina* actually constitute two sibling species that are not even closely related. The South American form is here described as a new species. The phylogenetic tree brings to light that *Girardia* arose on the South American portion of Gondwanaland, from which it, subsequently, dispersed to the Nearctic Region, probably more than once.

## INTRODUCTION

The genus *Girardia* comprises about 52 valid species, the natural distribution of which covers the Americas, from Southern Argentina and Chile to Southern Canada, albeit that in North America it is no longer the dominant type of freshwater planarian(Sluys et al. 2005). Furthermore, species of *Girardia* have been introduced into many other regions of the world (cf. Benítez-Álvarez et al submitted and references therein). For Australia, occurrence of introduced *G. tigrina* was established (Sluys et al., 1995 and references therein), apart from three presumed autochthonous species of *Girardia* (cf. Grant et al. 2006; Sluys & Kawakatsu 2001). However, recent molecular work (Grant, Sluys & Blair, unpublished) revealed that the latter three species (*G. sphincter* Kawakatsu & Sluys, 2001; *G. graffi* (Weiss, 1909); *G. informis* Sluys & Grant, 2006) do not belong to the genus *Girardia*.

Since the most recent, more comprehensive account on species of *Girardia* from the South American continent and the Caribbean Region by Sluys et al. (2005), 13 new species have been described (Chen et al. 2015; Souza et al. 2016; Souza et al. 2015; Hellmann et al. 2018, 2020; Lenguas Francavilla et al. 2021; Morais et al. 2021). Phylogenetic analyses of the genus *Girardia* are limited to the study of Sluys (2001), while historical biogeographic studies focusing on the genus are basically absent.

Due to great anatomical similarities between species, morphology-based phylogenetic analyses struggled to resolve the affinities between species and species-groups (cf. Sluys 2001). This problem is exacerbated by the fact that populations of *Girardia* may show only asexual reproduction by fissiparity and, thus, do not exhibit a copulatory apparatus, which hampers taxonomic identification, as well as the extraction of phylogenetic characters. Currently, the use of molecular markers allows overcoming some of the limitations of morphological characters to delimit species and to reconstruct the evolutionary history of triclads. The nuclear gene *Elongation Factor 1 alpha* (*EF1a*) has been used in several phylogenetic and phylogeographic studies in triclads (see Álvarez-Presas & Riutort, 2014), while the mitochondrial gene *Cytochrome Oxidase I* (*COI*) has been used for taxonomic studies as well as for species delimitation in the genus *Dugesia* (Sluys et al., 2013; Solà et al. 2015; Dols-Serrate et al. 2020; Leria et al., 2020). In all of these studies, an integrated approach, combining molecular and morphological data, proved to be highly successful in furthering our knowledge on the systematics of the various groups.

Here, we present the first molecular phylogeny of the genus *Girardia*, which resulted in several new insights into its taxonomic diversity, particularly in Mexican and South American territories, and into biogeographic history of the genus.

## MATERIALS AND METHODS

### Taxon sampling

For molecular analyses, samples of *Girardia* were obtained from Asia, Australia, Hawaii, the Americas, and Europe, with greater representation of the two last-mentioned geographical areas. Most of the samples were collected by the authors, while the rest was made available by various colleagues. Individuals were fixed in absolute ethanol. Specimens were identified to species level when both external and internal morphology could be examined (Table S1). When no anatomical information was available, individuals were simply classified as *Girardia* sp. In addition, all available *Girardia* sequences of *Cytochome Oxidase I* (*COI*) and *Elongation factor 1 alpha* (*EF1*_α_) were downloaded from GenBank. During our analyses some of the latter were excluded because of one or more of the following reasons: (a) low quality or short length of the sequences, (b) uncertain classification of the specimen, (c) avoidance of multiple sequences from a single locality (Table S2).

Figure 1 shows the distribution map of our samples, while an interactive map that allows a better resolution of the information on each locality is available at https://www.ub.edu/planarian-maps/. When available, we used the coordinates of the original samples, otherwise, approximated geographical coordinates were obtained from Google Earth (https://www.google.com/earth/index.html; last visited 12 March 2019) by entering the sampling localities (Table S1). We placed the data points on an open-source map (https://www.openstreetmap.org/) by using a custom script (not available in this publication) of JavaScript (ECMAScript 2015).

**Figure 1.**
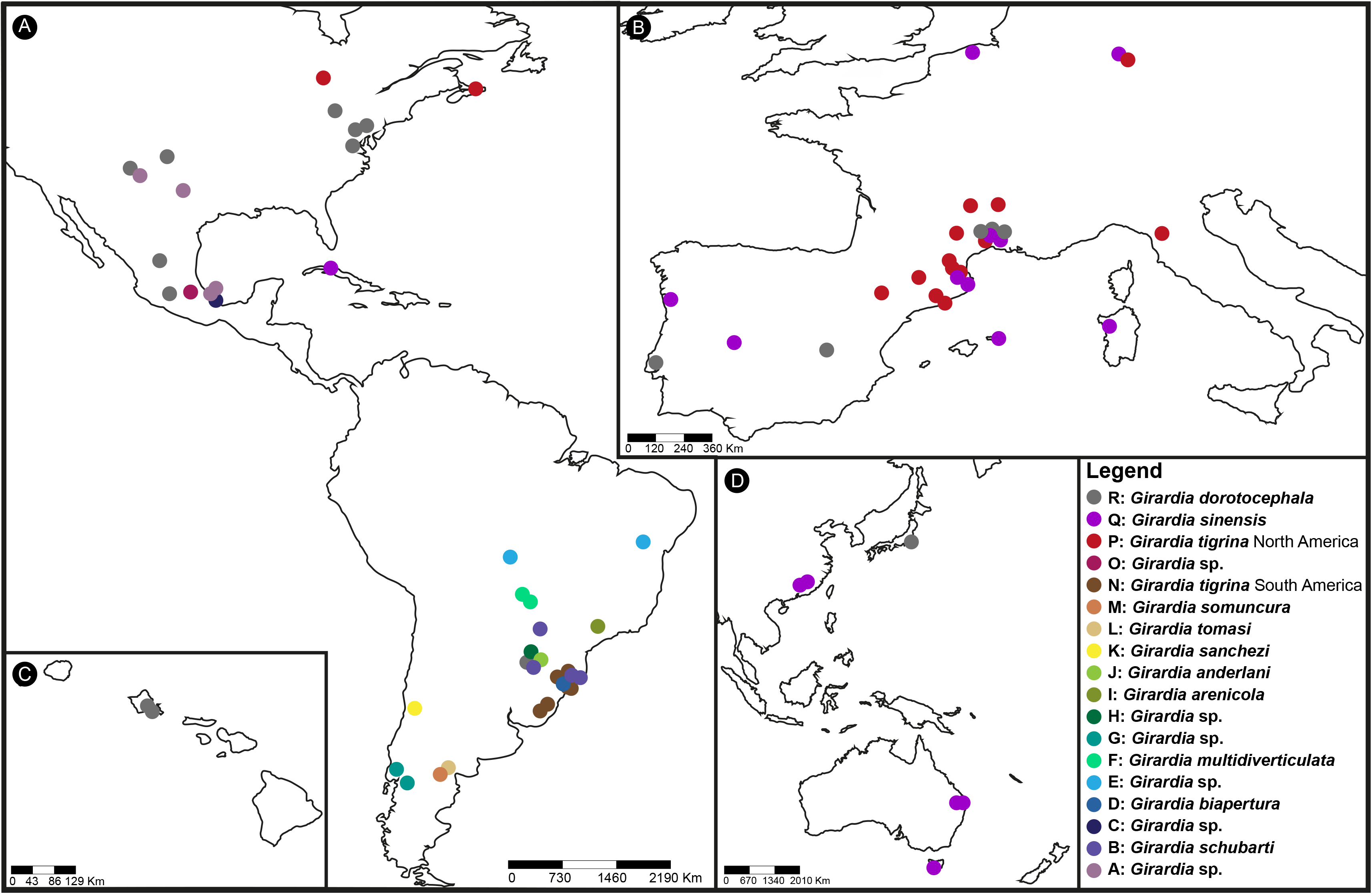
Maps showing the sampling localities for all individuals analyzed in the present study, including those corresponding to GenBank sequences. A: The Americas; B: Western Europe; C: Hawaii; D: Asia and Oceania. For a finer resolution, visit the interactive map at: https://www.ub.edu/planarian-maps/

### DNA extraction, gene amplification and sequencing

Genomic DNA was extracted using Wizard® Genomic DNA Purification Kit (Promega) and DNAzol® Reagent (Thermo Fisher Scientific, USA), according to the manufacturer’s instructions. The extraction product was quantified using a NanoDrop 2000c spectrophotometer (Thermo Fisher Scientific, USA).

A portion of the mitochondrial *COI* and of the nuclear *EF1*_α_ regions were amplified by Polymerase Chain Reaction (PCR), using 100 ng of template DNA and specific primers (Table 1) in 25µl of final reaction volume with MgCl_2_ (2.5mM), dNTPs (30μM), primers (0.4μM) and 0.75U of Go Taq® DNA polymerase enzyme (Promega Madison, Wisconsin, USA) with its buffer (1x). The amplification program consisted of 2 minutes (m) for initial denaturation at 95°C and 35 cycles of: 50 seconds (sec.) at 94°C, 45 sec. at annealing temperature (Table 1) and 50 sec. at 72°C; with a final extension step of 4 min. at 72°C.

**Table 1.**
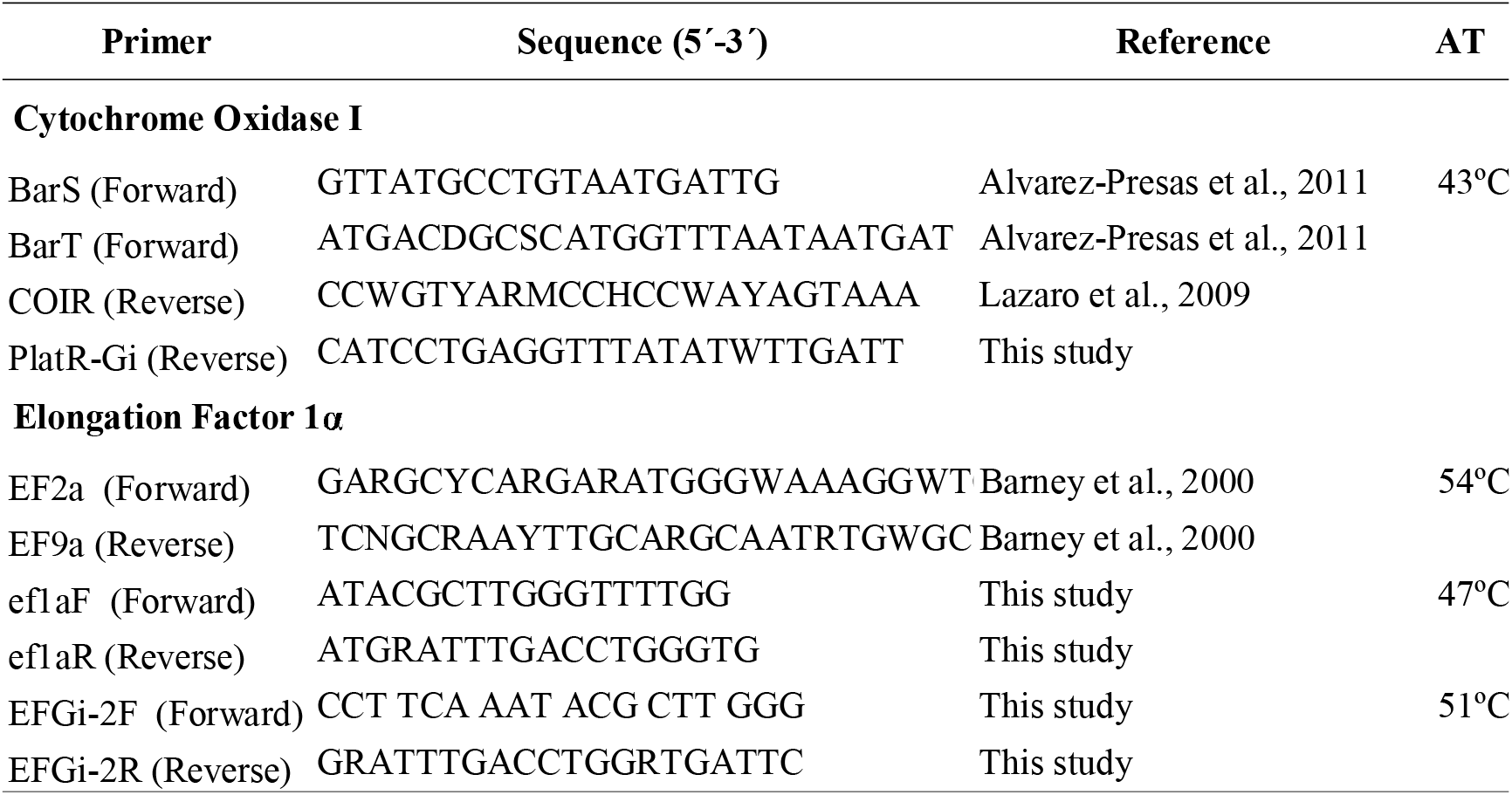
Primers used in this study, sequences, references, and annealing temperature (AT).

PCR products were run in agarose gels (1%) to check whether the correct band had been amplified. PCR products were purified by ultrafiltration in a Merck Millipore MultiScreen System (Darmstadt, Germany). For those samples that showed a faint PCR band on the electrophoresis, remaining PCR primers and dNTPs were digested by ExoSAP, a mix of two hydrolytic enzymes (Exonuclease I and Shrimp Alkaline Phosphatase; Thermo Fisher Scientific, USA) in a 3:1 ratio (amplified product: ExoSAP). Both strands of purified fragments were sequenced by Macrogen Inc., (Macrogen Europe, Madrid) with the same primers as used in the amplification. In order to obtain the final contigs, chromatograms were analysed with Genious v.10 (Kearse et al. 2012).

### Sequence alignment and datasets

Sequences of *COI* and *EF1*_α_, were aligned independently with ClustalW on the BioEdit Sequence Alignment Editor (Hall 1999). Each gene was translated into amino acids with the corresponding genetic code to check for the absence of stop codons and to produce the alignment, and, thereafter, converted again to nucleotides. Two alignments for each gene were generated, one including only *Girardia* sequences and the other comprising sequences of the closely related outgroup genera *Schmidtea* and *Dugesia* (cf. Álvarez-Presas & Riutort, 2014). We obtained the following four datasets for single gene alignments: (a) *COI* no outgroup (Dataset1), (b) *COI* with outgroup (Dataset2), (c) *EF1*_α_ no outgroup (Dataset3), (d) *EF1*_α_ with outgroup (Dataset4) (Table 2). The number of individuals sequenced for *COI* and *EF1*_α_ differ for the following reasons: (1) *COI* sequences were obtained first from nearly all samples, whereafter the phylogenetic tree was used to select samples for *EF1*_α_ sequencing by including individuals from different clades, as well as different localities; (2) for some samples, *COI* amplification was impossible and, thus, only *EF1*_α_ was obtained; (3) some species from GenBank have sequences for only one of the two markers (Appendix 1).

**Table 2.** Datasets analysed in this study, with their shorthand description, indication of the phylogenetic trees resulting from the analysis, number of species belonging to either *Dugesia* or *Schmidtea* used as outgroups, number of gene sequences, and the total length of these sequences in nucleotides.

| Data sets | Description | Tree figure | Outgroup |  | Gene sequences |  | OTUs | Length |
| --- | --- | --- | --- | --- | --- | --- | --- | --- |
|  |  |  | *Dugesia* | *Schmidtea* | COI | EF1a |  |  |
| Dataset1 | COI no outgroup | Figure S2 A |  |  | 124 |  | 124 | 837 |
| Dataset2 | COI with outgroup | Figure 2 | 6 | 3 | 133 |  | 133 | 840 |
| Dataset3 | EF1a no outgroup | Figure S2 B |  |  |  | 81 | 81 | 879 |
| Dataset4 | EF1a with outgroup | Figure S1 | 4 | 1 |  | 86 | 86 | 879 |
| Dataset5 | Concatenated no outgroup | Figure 3 |  |  | 94 | 78 | 98 | 1716 |

A concatenated dataset of both genes without outgroup (Dataset5) (Table 2) was obtained in Mesquite v3.04 (Massidon & Maddison, 2015), including all individuals for which sequences of both genes were available, as well as a few samples lacking one of the sequences. However, the latter sequences had to be included because they concerned the only available representatives for particular clades. Missing data were coded by Ns.

### Phylogenetic Inference

The best sequence evolution model and partition scheme for each gene alignment was estimated independently with PartitionFinder v2.1.1 (Lanfear et al. 2012), thereby considering the score for the Bayesian Information Criterion (BIC). As a preliminary step we hypothesized three partitions, corresponding with the first, second and third codon position for each gene. The results of the PartionFinder program validated this codon partition scheme, both for *COI* and *EF1*_α_. For each partition the best model was General Time Reversible + Gamma Distribution + Invariable Sites (GTR + Г + I). This codon partition scheme was then implemented in phylogenetic inference analyses, with the estimations of the parameters for each partition being independent.

Because nucleotide substitution saturation may decrease phylogenetic information contained in the sequences, a saturation test (Xia et al. 2003; Xia and Lemey 2009) was run, using DAMBE (Xia 2017). Third codon positions were analysed alone, while first and second positions were analysed together, including only fully resolved sites. Since the test can only analyse 32 Operational Taxonomic Units (OTUs) at a time, 10,000 replicates of subsets of 4, 8, 16, and 32 OTUs were performed. The proportion of invariant sites was calculated and included in the saturation analysis.

Bayesian Inference (BI) method was applied on the five data sets (Table 2) to infer the best tree and the posterior probabilities (PP), using MrBayes v3.2.2 (Ronquist et al. 2012). The chains were parameterized to 10 million generations, sampling every 1000 generations, and a 25% burn-in (default setting) was applied. Convergence of parameter values and topologies was examined by checking that the average standard deviation of split frequencies was below 0.01. Estimated sample size values (ESS) of each run were inspected in Tracer v1.5 (Rambaut et al. 2018) to check that the values were over 200.

## RESULTS

### Sequences and alignments

A total of 124 *Girardia* sequences of *COI* (109 obtained in this study, 15 downloaded from GenBank) and 81 of *EF1*_α_ (of which 80 new) are used in the final analyses, representing localities from all over the range of the genus (Appendix 1, Fig. 1). Several representatives of the genera *Dugesia* and *Schmidtea* are used as outgroups (Appendix 1). Sequences are analysed in individual gene alignments or combined into five datasets (Table 2). The alignments for each gene, including outgroups, show no saturation for any codon position, as determined by the saturation test (in all four tests Iss is significantly lower than Iss.c, thus indicating no saturation).

### Phylogenetic analyses

Phylogenetic trees were obtained from the five datasets (Figs 2, 3, S1, S2A, S2B). All phylogenetic trees delimit the same major clades and singletons (denoted with letters A to R in the trees), the composition of which does not change between datasets. These clades are fully supported (>0.99 PP) in the concatenated dataset, with the only exception of clade K (0.67 PP support). In the following, we will first describe the composition of the various clades and, where possible, the species assignments, followed by an account on the phylogenetic relationships between the clades.

**Figure 2.**
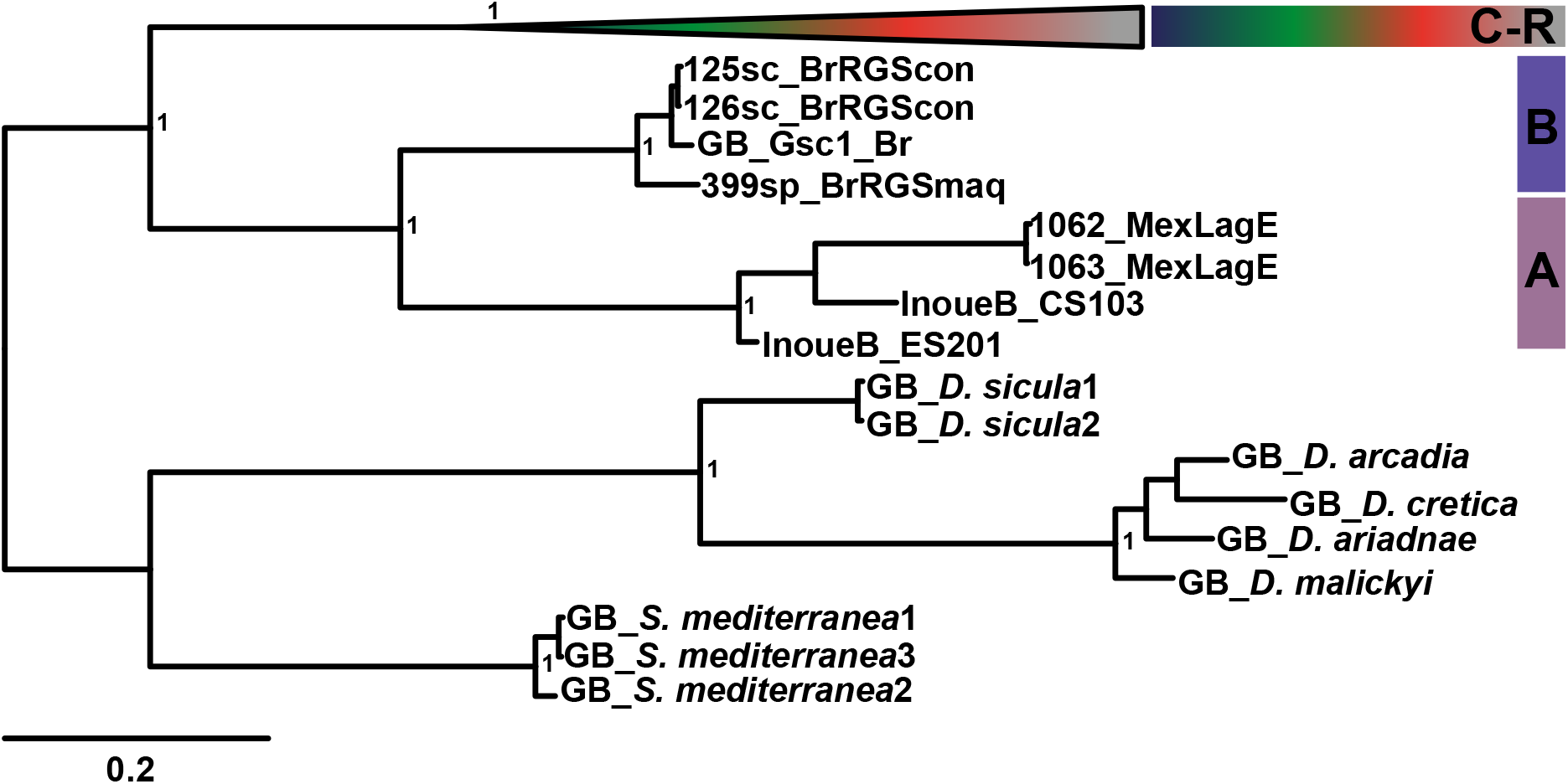
Bayesian Inference tree inferred from Dataset2 (*COI* with outgroup). Clades C to R have been collapsed for the sake of clarity. Clade A is constituted by unclassified samples from Mexico and Texas (USA); Clade B includes identified individuals of *Girardia schubarti* from Brazil and other unidentified Brazilian individuals. Outgroup is composed of several representatives of genera *Dugesia* and *Schmidtea* downloaded from GenBank (Appendix 1). Values at nodes correspond to posterior probability. Scale bar: number of substitutions per nucleotide position.

**Figure 3.**
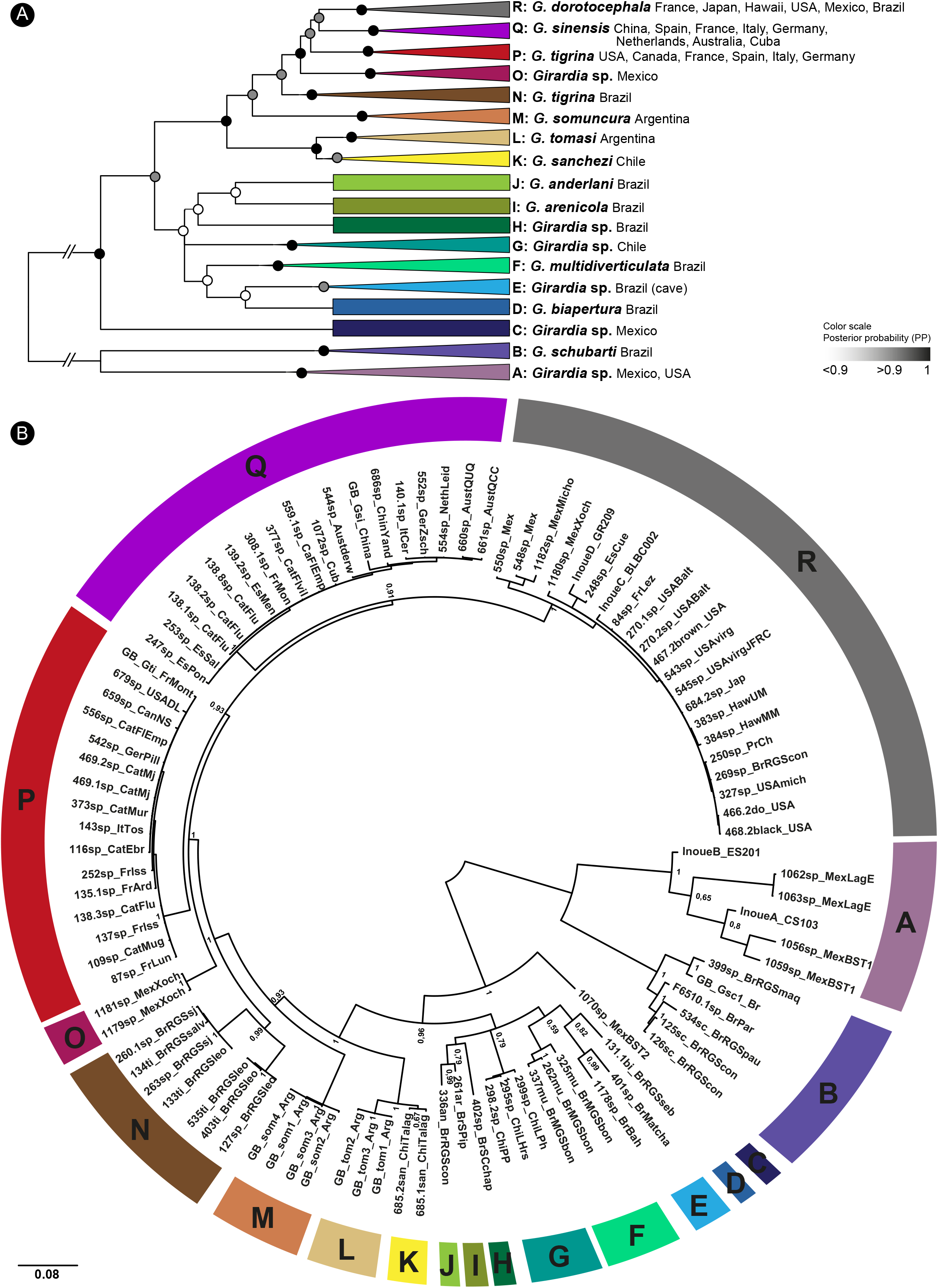
Bayesian Inference tree inferred from Dataset5 (concatenated no outgroup). Different clades indicated by letters and colours. (A) Schematic representation of the tree with collapsed clades, showing species identifications, when available, and countries of origin of the various terminals; shaded circles at nodes indicate posterior probability support, with values below 0.9 considered to be unsupported. (B) circular tree with all terminals; values at nodes correspond to posterior probability support. Scale bar: number of substitutions per nucleotide position.

#### Species assignment of the terminals and clades

Sequences of individuals of *G. schubarti* (Marcus, 1946) from GenBank and from two localities in southern Brazil constitute a monophyletic group, together with some unidentified specimens from two other localities in southern Brazil (clade B; Fig. 3A, B). This clade was highly differentiated from the rest of the OTUs in the tree, suggesting that it comprises, most likely, a single species, viz., *G. schubarti*. However, given its high diversity, this clade may actually correspond to a complex of species closely related to *G. schubarti*.

OTUs of *G. multidiverticulata* de Souza et al., 2015 (clade F), *G. biapertura* Sluys, 1997 (clade D), *G. anderlani* (Kawakatsu & Hauser, 1983) (clade J) and *G.* aff. *arenicola* Hellmann & Leal-Zanchet, 2018 (clade I) all group into their own clades, thus representing distinctly separated lineages (Fig. 3A, B). The branch of *G. sanchezi* (Hyman, 1959), represented by two individuals from the type locality in Chile, constitute clade K. Although this is the only clade with rather low support (0.67 PP), it is well-differentiated from all other OTUs, while it is not closely related to any of the other Chilean individuals that are included in our analyses and that together constitute clade G. OTUs of *G. tomasi* Lenguas Francavilla et al., 2021 and *G. somuncura* Lenguas Francavilla et al., 2021 from Argentina group in their respective clades L and M. Among our OTUs, there are five individuals from Brazil that had been identified as *G. tigrina*. Four of these individuals group into clade N, together with three non-identified individuals from Brazil, all from Rio Grande do Sul, thus suggesting that all of these OTUs belong to this species (Fig. 3A, B).

Although we were unable to include into our analyses any *G. tigrina* individual from North America, where the species is native, that was unequivocally identified as a member of this species, clade P comprises OTUs from Michigan, USA (679sp_USADL) and Nova Scotia, Canada (659sp_CanNS) (Fig. 3B). Furthermore, a *G. tigrina* specimen from France, of which sequences were downloaded from GenBank, also falls within clade P. Therefore, we assigned this entire clade to the species *G. tigrina*. All other OTUs in clade P come from outside of the autochthonous area of distribution of *G. tigrina* (Fig. 3A, B).

There are two other species of which taxonomically identified specimens were included in our study, viz., *G. sinensis* Chen & Wang, 2015 from China and *G. dorotocephala* (Woodworth, 1897) from North America. The first-mentioned species was represented by a sequence available from GenBank and the second by specimens purchased from Carolina Biological Supply Company and that were collected from the USA, albeit that exact provenance of this sample was not known. The sequences of these two species group into two separate clades, viz., clade Q (*G. sinensis*; GB_Gsi_China) and clade R (*G. dorotocephala*; 466.2do_USA) (Fig. 3B). However, apart from taxonomically identified *G. dorotocephala,* clade R houses also non-identified OTUs from USA, Mexico, Canada, Europe, Japan, Hawaii, and Brazil. Surprisingly, clade Q not only comprises taxonomically identified *G. sinensis* from China, but also non-identified OTUs from Australia, China, Cuba, and Europe.

Six clades (A, C, E, G, H, and O) can not be assigned to any known species of *Girardia*. Clade A comprises OTUs from very distant localities: Los Tuxtlas, Mexico (1062, 1063, 1056, 1059), and Texas (InoueA_CS103) and New Mexico (InoueB_ES201) in the USA. Clade C is constituted by an unclassified OTU from Los Tuxtlas, Mexico (1070), while clade E is formed by unclassified samples from two very distant (2018 km) Brazilian caves (401, 1178). Clade G comprises unclassified samples from Huinay Research Station, Chile (295, 298, 299), while clade H is formed by an unclassified sample (402) from Santa Catarina, Brazil. Clade O houses unclassified individuals (1181, 1179) from Xochimilco Mexico (Fig. 3B).

#### Phylogenetic relationships between clades

The phylogenetic trees based on the individual genes and including the outgroup taxa reveal the presence of two main, well-differentiated lineages within the genus *Girardia* (Figs 2, S1). One of these lineages, with OTUs from Mexico, USA, and Brazil, includes two sister clades (A+B), each of which is highly supported (1 PP), and that are well-separated from each other by long branches. The other main lineage comprises all remaining *Girardia* samples, with OTUs from North, Central and South America, which group in the clades C-R (these clades are collapsed in Figs 2 and S1). Three of the lineages in this second main group (clades P, Q, R), concern OTUs that have been introduced into other parts of the world, outside of the native range of *Girardia*.

In view of these results, we replaced in further phylogenetic analyses the initial outgroup taxa (species of *Dugesia* and *Schmidtea*) by the A+B clade, in order to avoid rooting with outgroup taxa that might be too distantly related to the ingroup. In this way, we attempted to avoid long-branch attraction (Felsenstein 1978) and systematic error due to highly divergent outgroup taxa (Graham et al. 2002).

Phylogenetic trees resulting from analyses of both concatenated and individual-gene datasets, rooted with clade A+B, are shown in Figure 3 and Figure S2. Individual-gene analyses (Fig. S2) recovered less internal nodes that are fully supported than the analysis of the concatenated dataset (Fig. 3), probably due to synergetic information in the molecular markers. In the following we describe the relationships and supports found in the concatenated tree (Fig. 3).

With respect to the ingroup C-R, an unclassified OTU from Los Tuxtlas (clade C) is sister to a major branch comprising all remaining clades, with good support (0.96 PP). One branch of this major clade comprises the groups D-J, which does not receive high support (0.79 PP), and concerns several South American lineages with unresolved affinities, such as: *G. biapertura* (D); OTUs from two Brazilian caves (E), and from Santa Catarina (H); *G. anderlani* (J) and the troglobitic *G.* aff. *arenicola* (I); the troglobitic *G. multidiverticulata* (F), and OTUs from Chile (G).

The second major branch on the tree, comprising clades K-R is highly supported (1.0 PP) (Fig. 3A, B). It contains a clade formed by the two sister species *G. sanchezi* (K) and *G. tomasi* (L), as well as clades of the following six well-differentiated taxa: *G. somuncura* (M), *G. tigrina* from Brazil (N), unclassified OTUs from Xochimilco, Mexico (O), *G. tigrina* from North America (P), *G. sinensis* (Q), and *G. dorotocephala* (R). All nodes within the K □ R group receive high to maximum support values, ranging between 0.91 and 1.0. Hence, the topology of this portion of the tree (Fig. 3) shows well-supported relationships, in contrast to clade D – J.

#### Historical biogeographic analysis

Since our schematic tree (Fig. 3A) includes the countries of origin of the various terminals, it allows for an historical biogeographic analysis of the genus *Girardia*, at least for the taxa included in our study. In this analysis, it is important to take into account a number of issues that complicate geographic interpretation of the tree.

For example, *G. sinensis*, although described from China, has a North American origin (see discussion). Moreover, *G. sinensis, G. tigrina*, and *G. dorotocephala* have been introduced from North America into other parts of the world, and, therefore, any country outside of the North American subcontinent should be disregarded in the analysis. However, South America is an exception to this rule, in that the present study shows that presumed *G. tigrina* from this subcontinent actually concerns the new, sibling species *G. clandistina* Sluys & Benítez-Álvarez, sp. nov. (see below). The introduction of species of *Girardia* from their native areas to other parts of the world, and their subsequent settlement and further dispersal, have been analysed in extenso by Benítez-Álvarez et al. (personal communication) and, therefore, shall not be discussed here any further.

From that perspective, it is clear that the ancestral distribution of the clade O-R concerns the North American subcontinent and that of clade D-J concerns the South American subcontinent. When the South American distributions of the clades K-L, M, and N are taken into account, it leaves little doubt that the ancestral distribution on the branch leading to clades D-R must be reconstructed as being South America, as, most likely, is the case also for the most basal branch, leading to all *Girardia* terminals included in the tree. In other words, the ancestral distribution of *Girardia* is South America, while the North American clade is the result of more recent colonization events.

## DISCUSSION

### *Girardia*: genetical and chromosomal divergences

Our phylogenetic tree indicated the existence of two major lineages of *Girardia*, one constituted by the sister taxa *G. schubarti* (clade B) and the taxonomically unidentified clade A, and the other comprising all other *Girardia* OTUs and taxa (C □ R) (Fig. 3). In a morphological phylogenetic analysis, *G. schubarti* grouped well among the other species of *Girardia* and formed a clade together with *G. arizonensis* (Kenk, 1975) and *G. azteca* (Benazzi & Giannini, 1971) (Sluys 2001). In their study on the phylogeny of continenticolan planarians with the help of molecular markers, Álvarez-Presas & Riutort (2014) included also three species of *Girardia* and found a sistergroup relationship between *G. anderlani* and *G. tigrina*, which together were sister to *G. schubarti*. The analysis of Inoue et al (2020) showed a sister-group relationship between *G. tigrina* and *G. dorotocephala*, which together with *G. anderlani* and two putative new species (InoueC and InoueD in our trees) constituted a monophyletic group that was sister to a clade formed by *G. schubarti* and two other putative new species (InoueA and InoueB in our trees). Lázaro et al. (2011) found also a sister-group relationship between *G. tigrina* and *G. dorotocephala*, which together were sister to *G. schubarti*, being the only three species of *Girardia* included in their analysis. However, our phylogenetic tree indicates a sister-group relationship between *G. schubarti* (clade B) and the taxonomically unidentified clade A (including InoueA and InoueB species), both of these clades together being sister to the major branch comprising all other *Girardia* OTUs and taxa (C □ R) in our analysis. Evidently, as our study includes more OTUs than those of Álvarez-Presas & Riutort (2014) and Inoue et al. (2020), a more complex pattern of genealogical affinities is to be expected. In addition to the clear sister-group relationship between clades A+B and C □ R, the great genetic distance between these two clades is noteworthy (Fig. 2), since the length of the branches is comparable with the distance between the two sister genera *Schmidtea* and *Dugesia*, which presumably diverged about 135.9 million years ago (Mya) (Solà et al 2022).

In addition to this high genetic differentiation, *G. schubarti* is also differentiated from other *Girardia* species by its number of chromosomes, having a basic haploid complement of n=4 (Kawakatsu et al. 1984; Jorge et al. 2000; Knakievicz et al. 2007), albeit that similar chromosome portraits are found in *G. arizonensis* Kenk, 1975 and *G. jenkinsae* Benazzi & Gourbault, 1977 (Benazzi & Gourbault, 1977; Benazzi, 1982), species not included in the present study (unless they are represented by some of our unidentified specimens from Mexico or the USA). On the other hand, *G. tigrina*, *G. dorotocephala, G. sanchezi, G. anceps*, *G. tahitiensis*, and *G. festae* exhibit haploid complements of n=8 (Gourbault 1977; Puccinelli & Deri 1991), while *G. anderlani*, *G. biapertura* and *G. cubana* (Codreanu & Balcesco, 1973) have n=9 (Benazzi 1982; Jorge et al. 2000; Benya et al. 2007; Knakievicz et al. 2007). For *G. nonatoi,* Marcus (1946) counted in oocytes 10 chromosomes during meiosis in the haploid phase, so that the full complement presumably consists of 20 chromosomes. Unfortunately, the chromosome portraits of other species of *Girardia* are unknown. Despite this paucity of information on chromosome number in the genus *Girardia*, a pattern emerges when the complements are plotted on the phylogenetic tree: clade A+B includes *G. schubarti* with n=4, D □ J includes two species (*G. anderlani*, *G. biapertura*) with n=9, and K □ R clade includes three species (*G. tigrina*, *G. dorotocephala, G. sanchezi*) with n =8. If the chromosomal numbers found in the few species within each of these major clades (A+B, D-J, and K-R) are presumed to be common for all species within each of these groups, it may be hypothesized that the origin of the main clades of *Girardia* was associated with events of genomic duplications and/or chromosomal rearrangements.

Differences in chromosome number between closely related species of triclads are relatively common and have been related to speciation events in the genera *Schmidtea* and *Dugesia* (Leria, et al. 2018, 2020). However, with the present information available for *Girardia*, it cannot be excluded that chromosomal changes were not the drivers of the speciation process, but accumulated only after speciation had taken place. Therefore, it is only through future, more comprehensive and integrative studies that we may determine whether the great genetical and chromosomal divergences of the A+B clade, as compared to its congeners, warrant taxonomic recognition in the form of a separate genus, or merely represent highly evolved autapomorphic features for a particular branch within the genus *Girardia*.

### Genetic differentiation within clades of *Girardia*

Although we only have scattered samples from all over the Americas (Appendix 1, Fig. 1), in many cases our molecular-based phylogenetic results revealed a high genetic diversity and structure within *Girardia*, particularly in Mexico and Brazil.

Mexico showed the highest molecular diversity, despite the rather low number of samples. Eleven individuals were analysed, two of unknown origin and the rest coming from five localities (Appendix 1), which exhibited clear structure and genetic differentiation, and comprised four different clades (A, C, O, and R, Fig. 3). From these four clades, particularly clade A is noteworthy because of its high internal diversity, albeit that it does not include any taxonomically identified species. However, genetic structure within clade A strongly suggests that it contains more than one species. This is in accordance with the suggestion made by Inoue et al. (2020), who delimited two putative species on the basis of short fragments of *COI*, coded InoueA and InoueB in the present work (Fig. 3). Clades C and O (sister to D-R clade and P+Q+R clade, respectively) comprise only animals from Mexico, while some Mexican individuals occur also in clade R (*G. dorotocephala*). Evidently, at this moment it remains undecided whether the observed genetic diversity concerns new species of *Girardia* or merely reflects the presence of already known species of which molecular data is still lacking.

This high genetic Mexican diversity is not geographically structured as, perhaps, might be expected. Within clade A, the long branch separating samples from the Biological Station (1056, 1059) and individuals from Laguna Escondida (1062, 1063) suggests that two genetically highly differentiated species are present at these two localities, although the collection sites are only 2 Km apart. Clade C is only formed by individual 1070 from a second collection site at the Biological Station. All of this points to a possible co-occurrence of two highly differentiated species in the same river within the Biological Station. At the Xochimilco locality we found three specimens, two constituting the sister clade (O) of the group including clades P, Q and R, while the third specimen (1180) belongs to clade R (*G. dorotocephala*). This mix of genetically distant species at sites that geographically are in close proximity to each other, suggests a complex history for the diversification and evolution of *Girardia* in the Americas.

Another interesting fact that surfaced in our analyses was the relatively high diversity of cave-dwelling species in Brazil, with *G. multidiverticulata* being the first troglobitic continenticolan to be reported from South America. Although its distinctive characters differ from other species of *Girardia*, it shares with *G. anderlani* the presence of a large and branched bulbar cavity (Souza et al., 2015). Unfortunately, our molecular trees did not show sufficient resolution to support a sister-group relationship between *G. multidiverticulata* and *G. anderlani.* In point of fact, the trees suggested a closer relationship between epigean *G. anderlani* and specimen 261, which probably represents troglobitic *G. aff. arenicola*, both showing dorsal testes and a branched bulbar cavity (Hellmann et al. 2018).

Among our Brazilian samples there are two others that originated from hypogean habitats, viz., OTUs 401 and 1178, together constituting clade E (Fig. 3). Although these two individuals constituted a monophyletic group, they are genetically quite distinct, while their sampling localities are far apart. This suggests that clade E comprises two new cave-dwelling species of *Girardia*. This recently discovered flourishing of hypogean *Girardia* species in Brazil (see Morais et al. 2021) may be an indication that the genus is highly successful in adapting to life in caves and that future studies of those habitats in other regions in the Americas may unveil further diversity.

To date, nine species of *Girardia* have been recorded from Mexico, USA and Canada, with *G. tigrina* and *G. dorotocephala* being the most widely distributed ones (Sluys et al. 2005). The present study adds *G. sinensis*, described from a locality in China (Chen et al. 2015), since we identified it molecularly from Cuba (Figs 1, 3A). Moreover, the close phylogenetic relationship that this species shares with *G. dorotocephala* and *G. tigrina*, both of North American origin, also clearly point to that region as the original area of distribution of *G. sinensis*. In a recent molecular study on freshwater planarians from New Mexico and Texas, Inoue et al. (2020) identified two putative new species (InoueC and InoueD in our trees) that were closely related to *G. tigrina* and *G. dorotocephala*. However, in our analyses, both of their sequences grouped among representatives of *G. dorotocephala* (Fig. 3). Several other sequences from individuals collected outside of the Americas fall into the three clades P, Q, R, thus corroborating the North American origin of the introduced populations. The expansion of these three lineages around the world and its possible impact, have been analysed by Benítez-Álvarez et al. (submitted).

### Nominal Girardia tigrina

A very interesting result of our molecular analysis concerns the positions in the phylogenetic tree of the North and South American *G. tigrina* samples, showing that the Brazilian clade (N) is not even closely related to the North American one (clade P) (Fig. 3). This corroborates the conclusion of Sluys et al. (2005) that the North and South American forms are different species. According to Sluys et al. (2005), the only anatomical difference between them resides in the coat of muscles around the bursal canal. In North American *G. tigrina* this coat of muscles is simple, consisting of a thin subepithelial layer of circular muscle, followed by an equally thin layer of longitudinal muscle fibres. In contrast, the South American form possesses a bursal canal musculature that consists of a well-developed coat of intermingled circular and longitudinal muscle fibres. In other characters the two forms are very similar, but our results clearly show that they are genetically well-differentiated and are not even sister species. Therefore, the South American form is here described and named as the new species *Girardia clandistina* Sluys & Benítez-Álvarez, sp. nov. (for differential diagnosis, see Appendix 2). This taxonomic action is not unimportant, since *G. tigrina* is the type species for the genus *Girardia* (Kenk 1974) and, therefore, it is necessary to know the precise boundaries of the taxon and the extension of the species name.

### Historical biogeography of *Girardia*

Our phylogenetic tree suggested that *Girardia* evolved on the South American subcontinent and from there colonized North America. Previous studies argued that the family Dugesiidae, including *Girardia*, had already diversified on Gondwanaland (Ball 1975) or even at pre-Pangaean times and, thus, must have diversified also already on Pangaea (Sluys et al. 1998). Thus, *Girardia* diversified on the South American portion of Gondwanaland, and, subsequently, ancestors of the O-R clade migrated to North America and diversified there. The relatively short inter-branches among the clades on the North American subcontinent (clades O, P, Q, R; Fig. 3B) implies that diversification of *Girardia* in North America is rather recent, and hence that the northward migration for this group did probably take place only after complete closure of the Isthmus of Panama at about 2.8 Mya (O’Dea et al. 2016).

Interestingly, presence of the basal lineages A in Mexico and the USA, and clade C in Mexico, and of the crown group O-R in Mexico, USA, and Canada, suggests that the North American subcontinent was populated by at least two independent waves of dispersal from the Neotropics. Although in these cases, the available data do not allow us to infer whether this northward migration was relatively recent or took place in more remote epochs, again, it would have been possible only through freshwater tracks in the intermittent connections during emergence of the Isthmus of Panama or once it was fully established (McGirr et al. 2021). Unfortunately, lack of OTUs from the northern parts of South America prevents further elucidation of the precise routes taken by neotropical *Girardia*’s during their dispersal into the Nearctics.

## Acknowledgments

We thank all colleagues mentioned in Table S1 for contributing samples of worms. We are grateful to Oleguer Castillo, Ana Paula Goulart Araujo, and Omar Lagunas for DNA extraction and sequencing of some samples, while the first-mentioned also designed a specific *Girardia COI* primer. We are also grateful to Eduard Solà for designing primers for EF1a amplification. Alejandro Oceguera and Fernando Carbayo are thanked for support in the sampling campaign in Mexico.

## Data availability

All sequences have been deposited in GenBank.

## LEGENDS TO FIGURES, TABLES, SUPPLEMENTARY FIGURES, AND APPENDICES

**Appendix 1.** Sequences included in this study, with indication of sample codes, sampling localities (for *Girardia* samples), taxonomic assignment before and after analyses, codes used in the text and figures, and GenBank accession numbers for *COI* and *EF1*_α_ sequences. Sequence codes in bold concern new sequences reported in this study. See Supplementary Table S1 for exact localities, collectors, and criteria used for taxonomic assignment previously to our analysis.

**Appendix 2.** Differential diagnosis of *Girardia clandistina* Sluys & Benítez-Álvarez, sp. nov.

## SUPPORTING INFORMATION

Figure S1. Bayesian Inference tree from Dataset4 (EF1a with outgroup). The sister group of *Girardia schubarti* and unclassified samples from Mexico have been collapsed. Values at nodes correspond to posterior probability support. Scale bar: number of substitutions per nucleotide position.

Figure S2. Bayesian Inference trees from Dataset1 (*COI* without outgroup) (A) and Dataset3 (*EF1*_α_ without outgroup) (B). Values at nodes correspond to posterior probability support. Scale bar: number of substitutions per nucleotide position.

Table S1. Samples included in this study, geographical coordinates, collectors, and identification method. The asterisk indicates approximated coordinates based on sampling locality description.

Table S2. *COI* and *EF1*_α_ sequences of *Girardia* genus present in GenBank. In bold are indicated the sequences included in the analyses. Reason for exclusion is indicated for certain sequences not included in the present study.

## Appendix 1

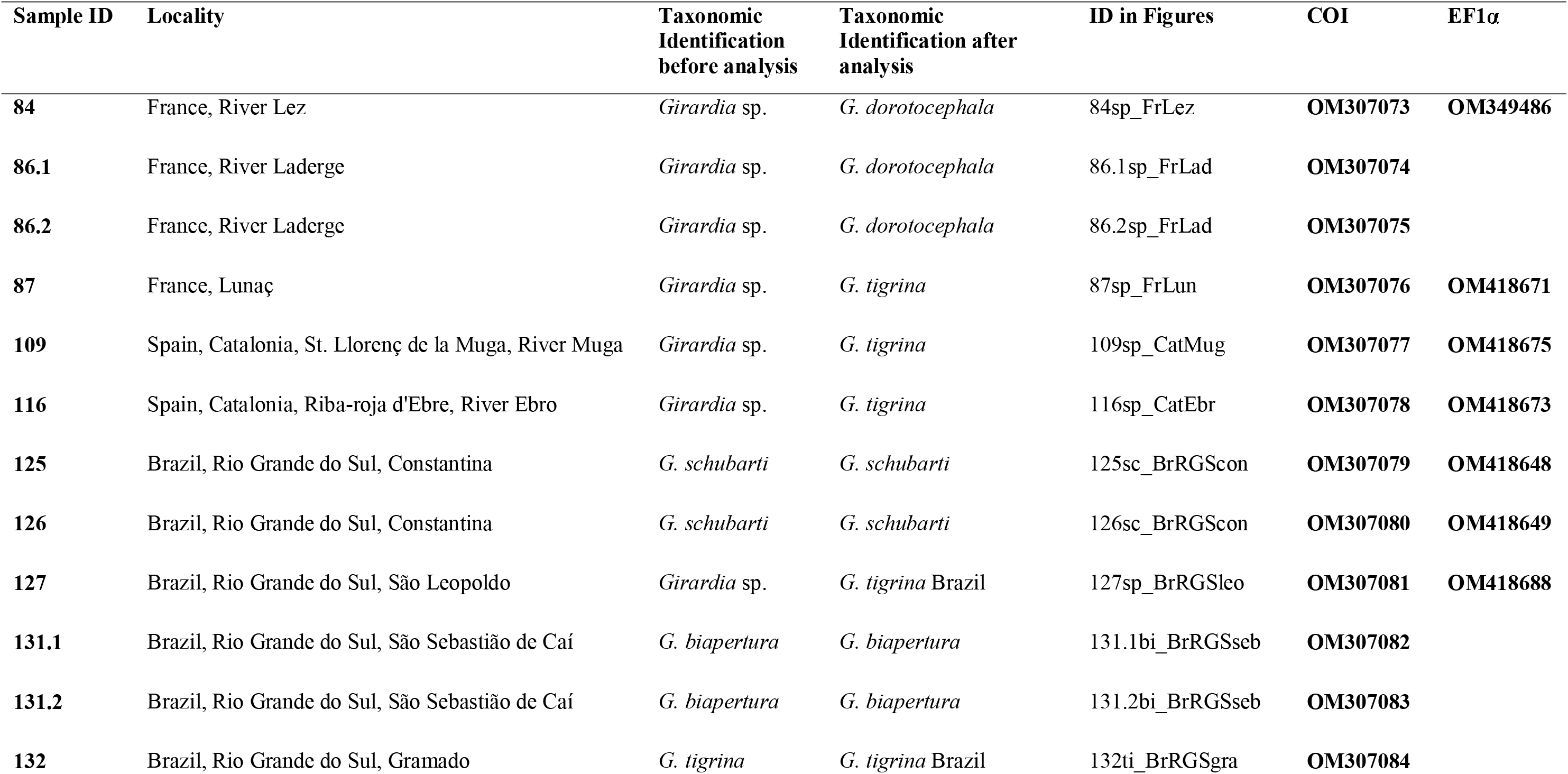
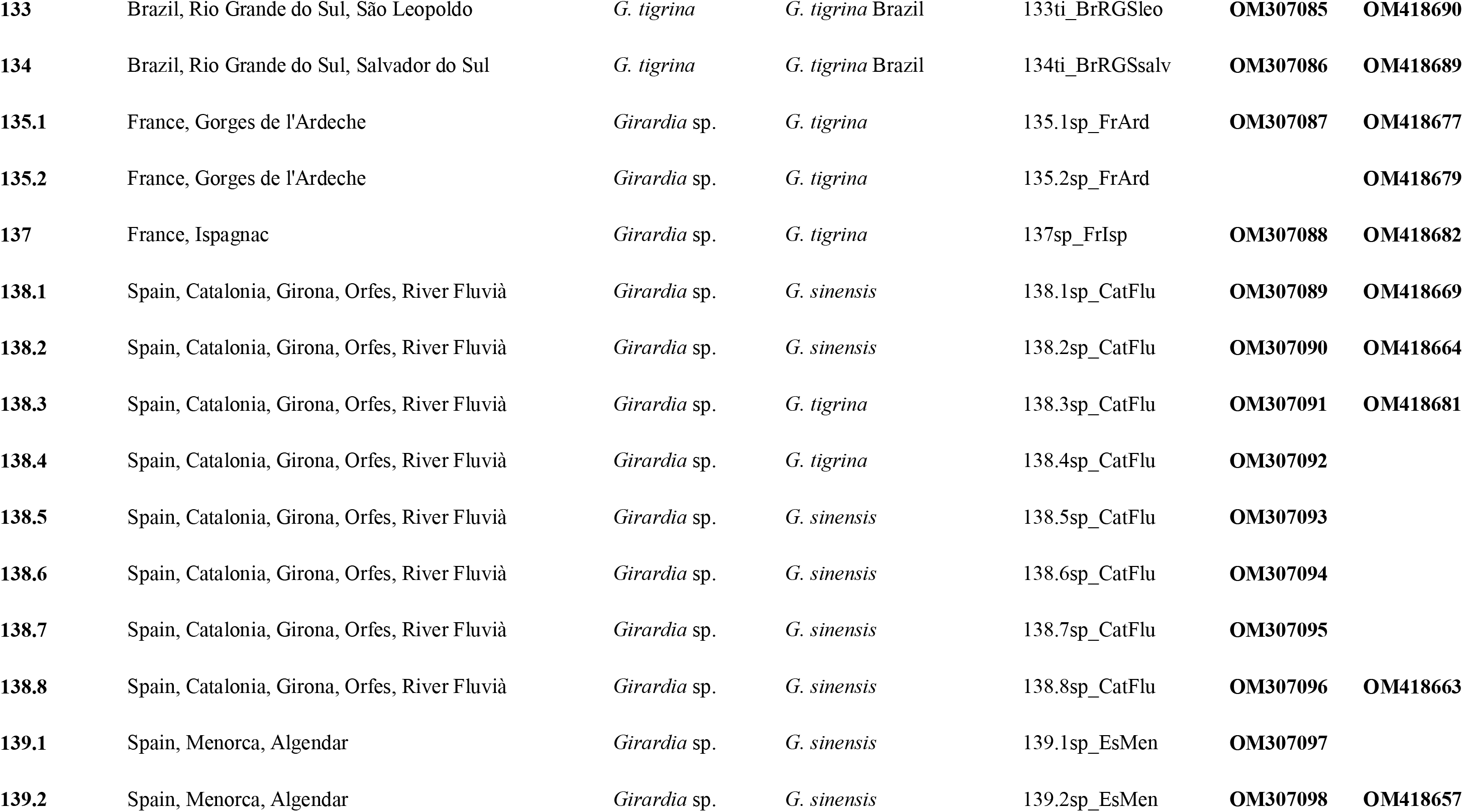
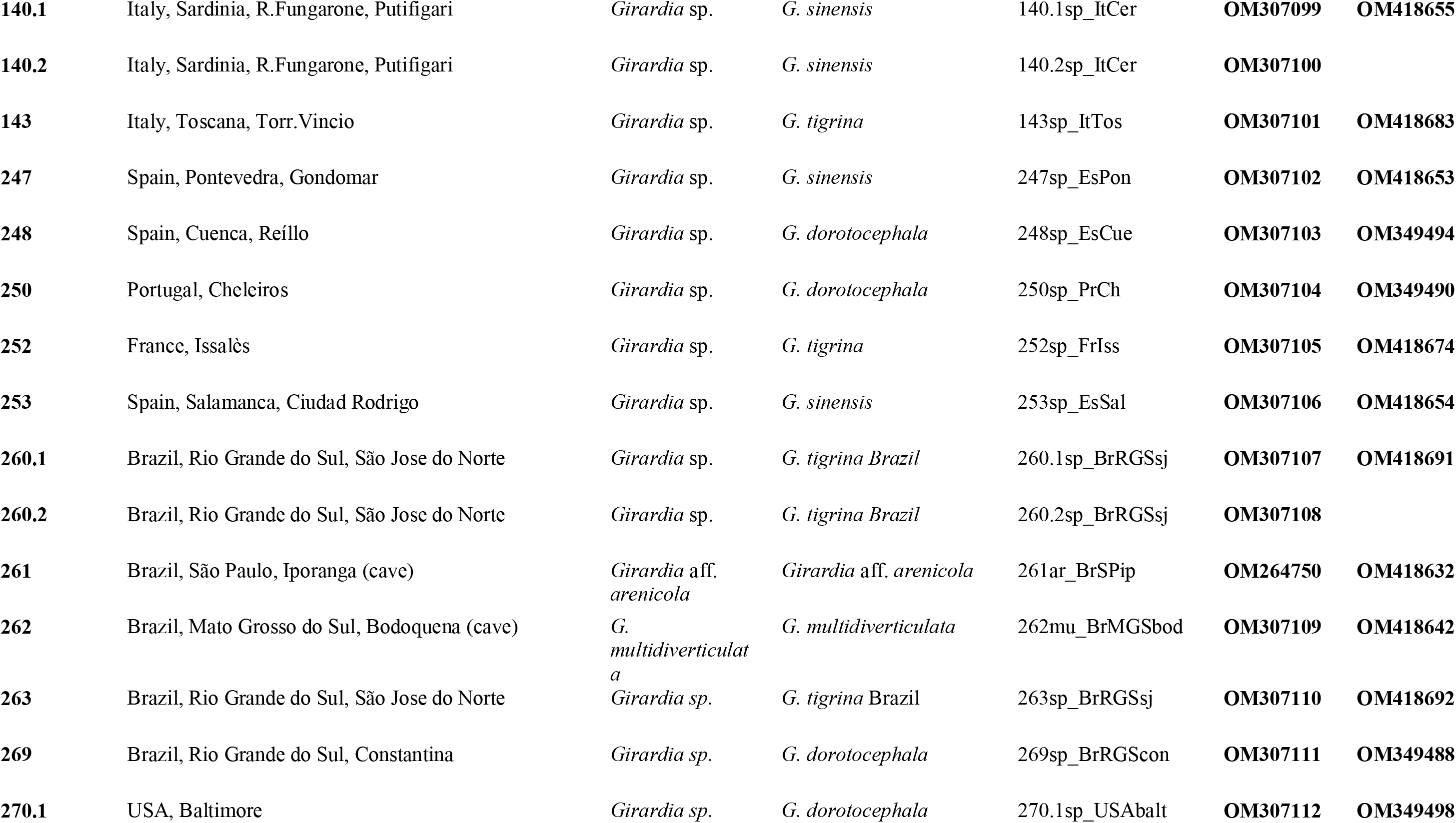
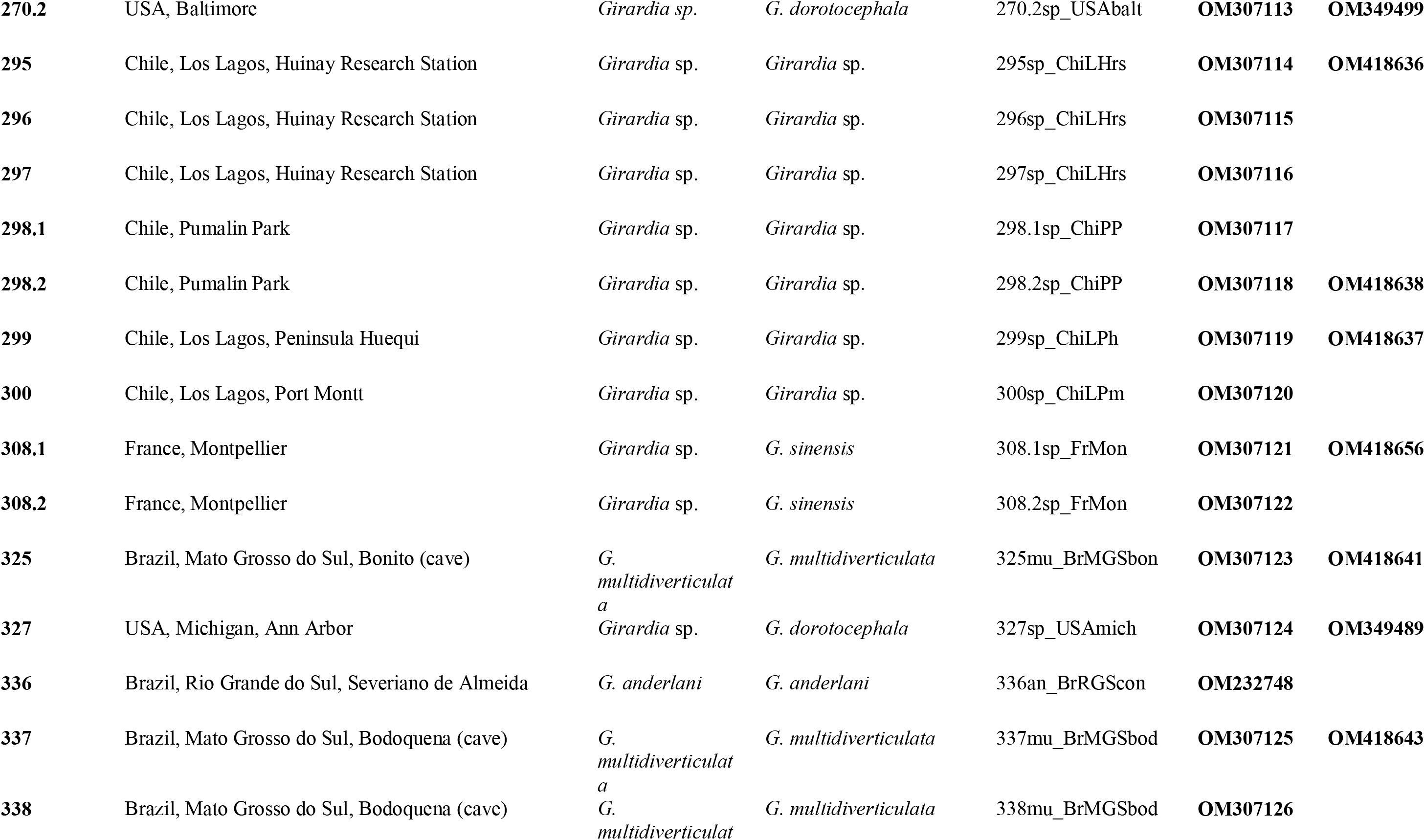
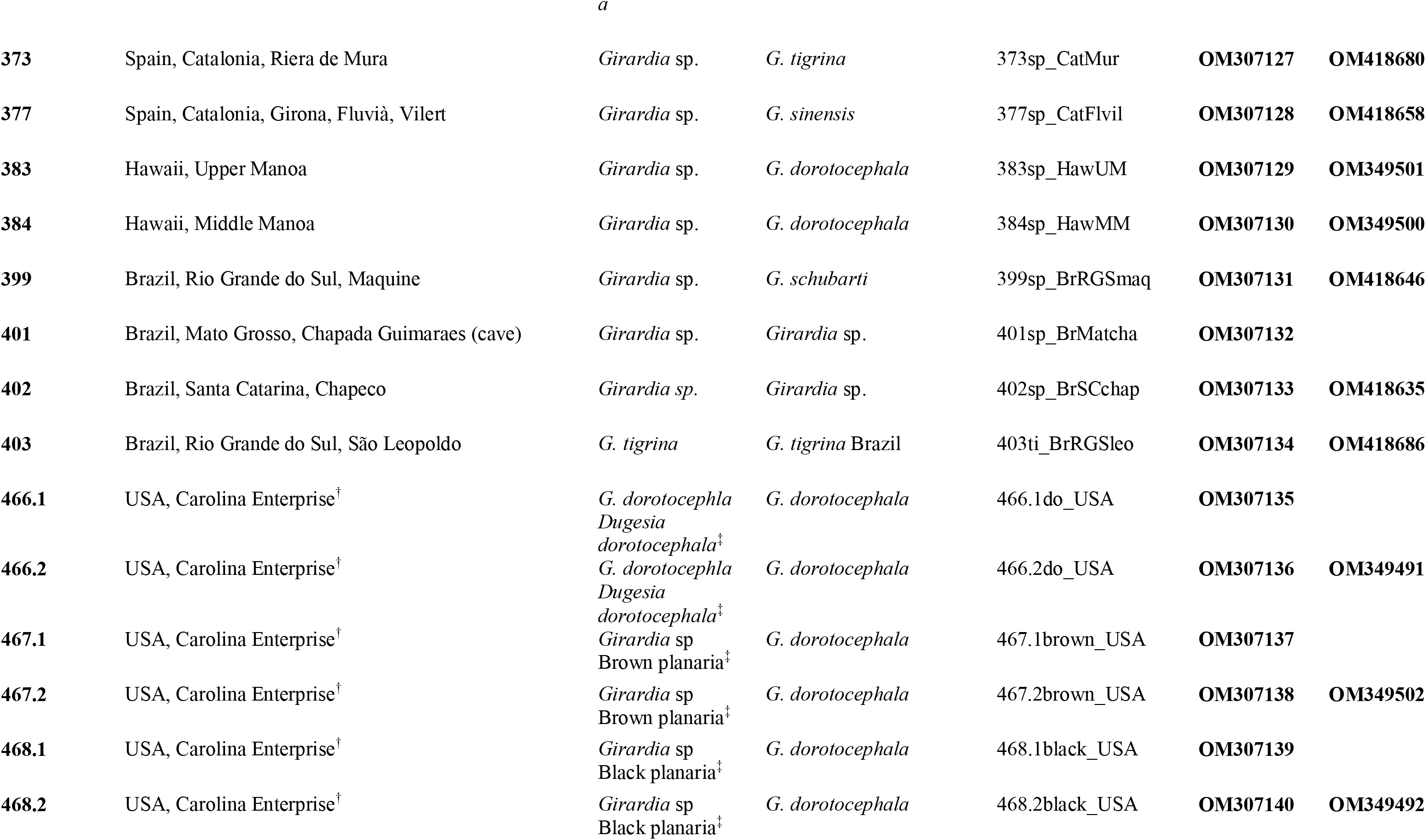
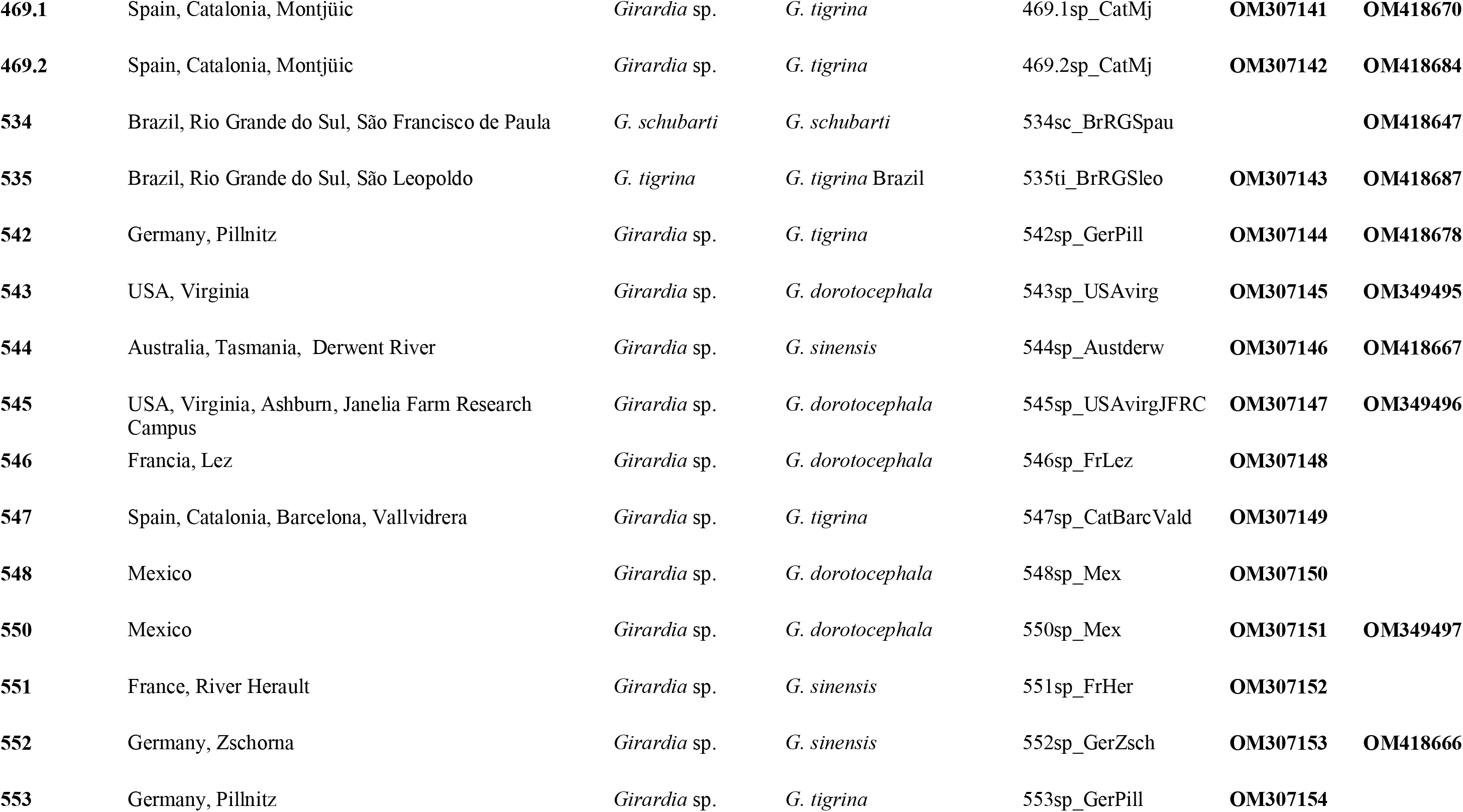
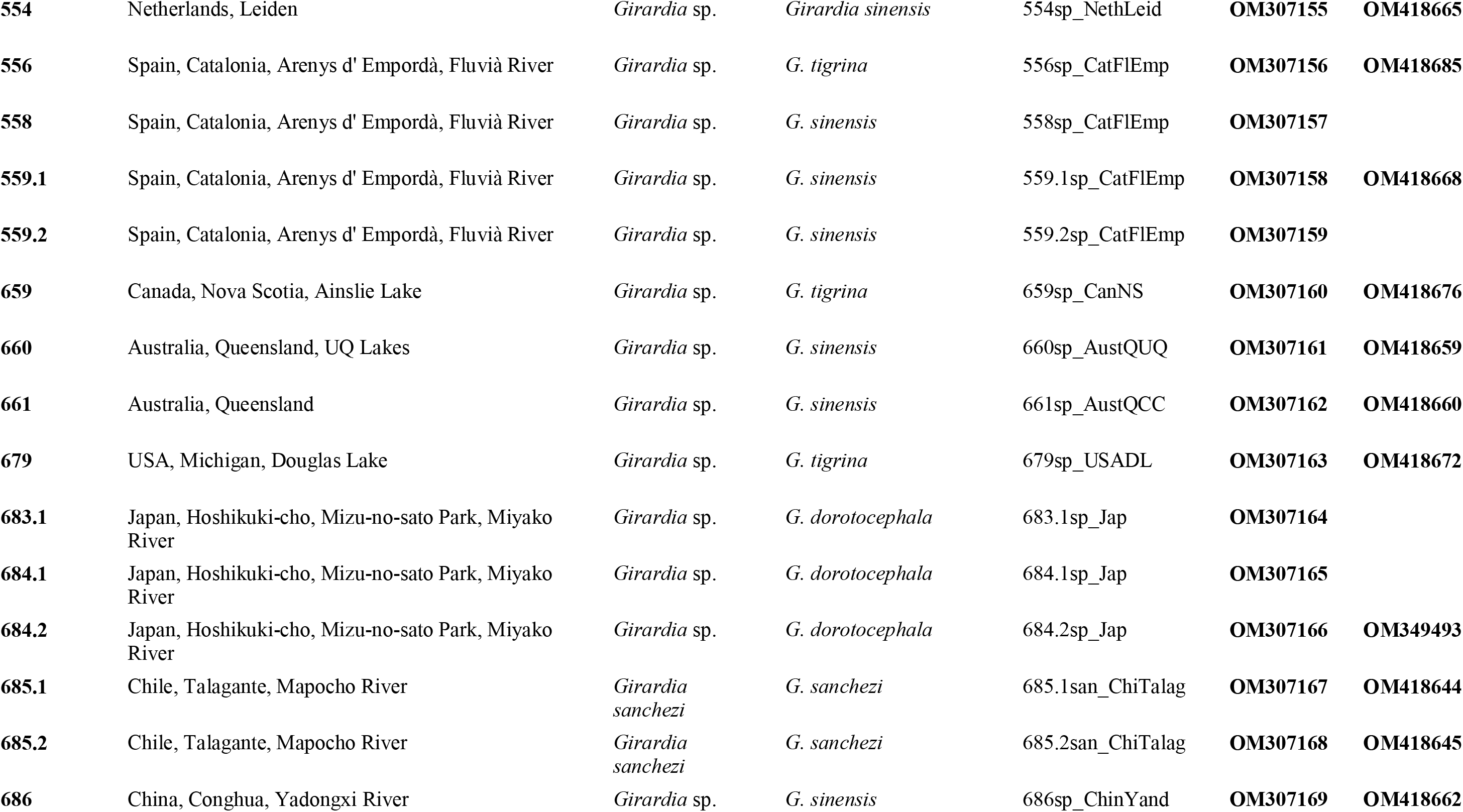
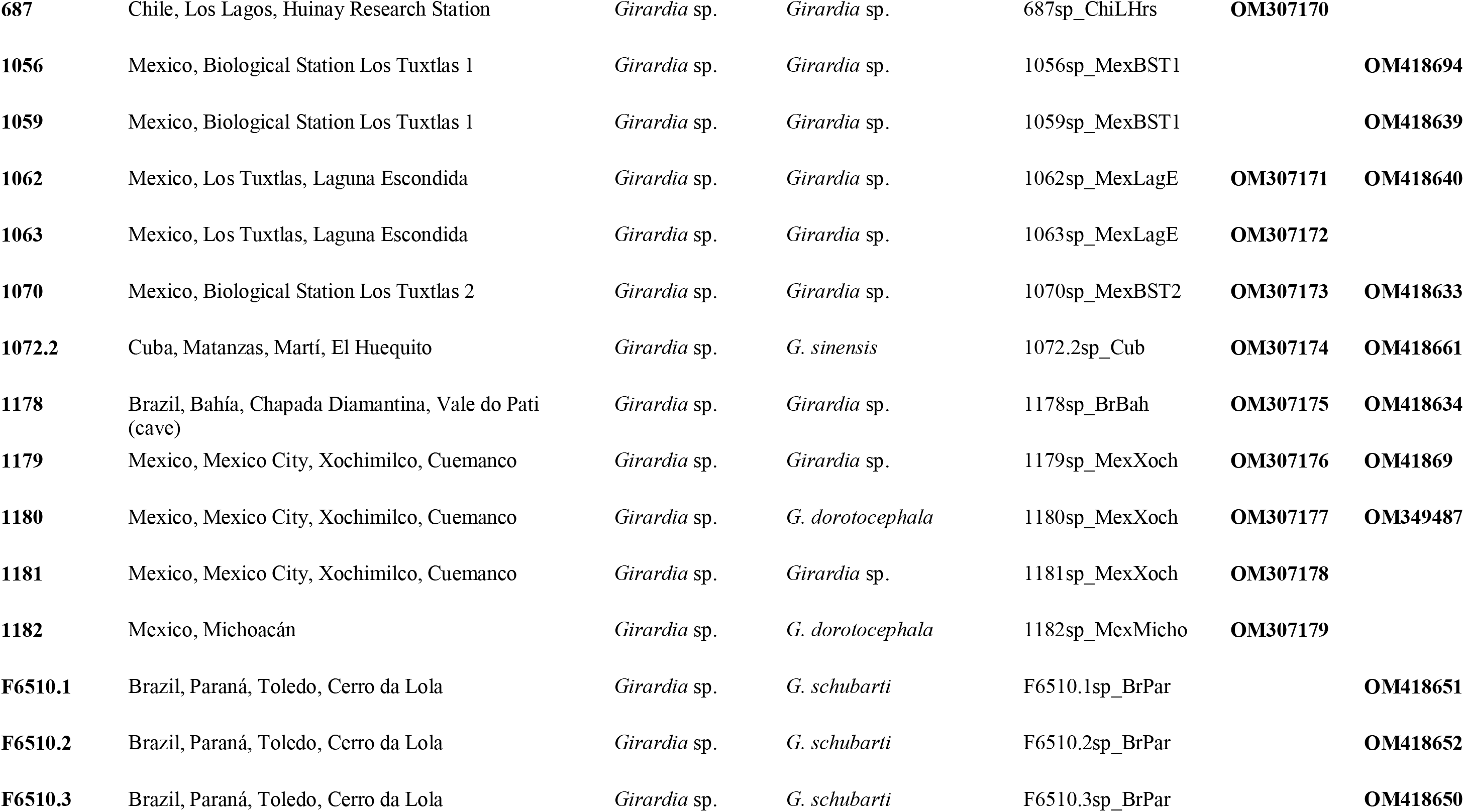
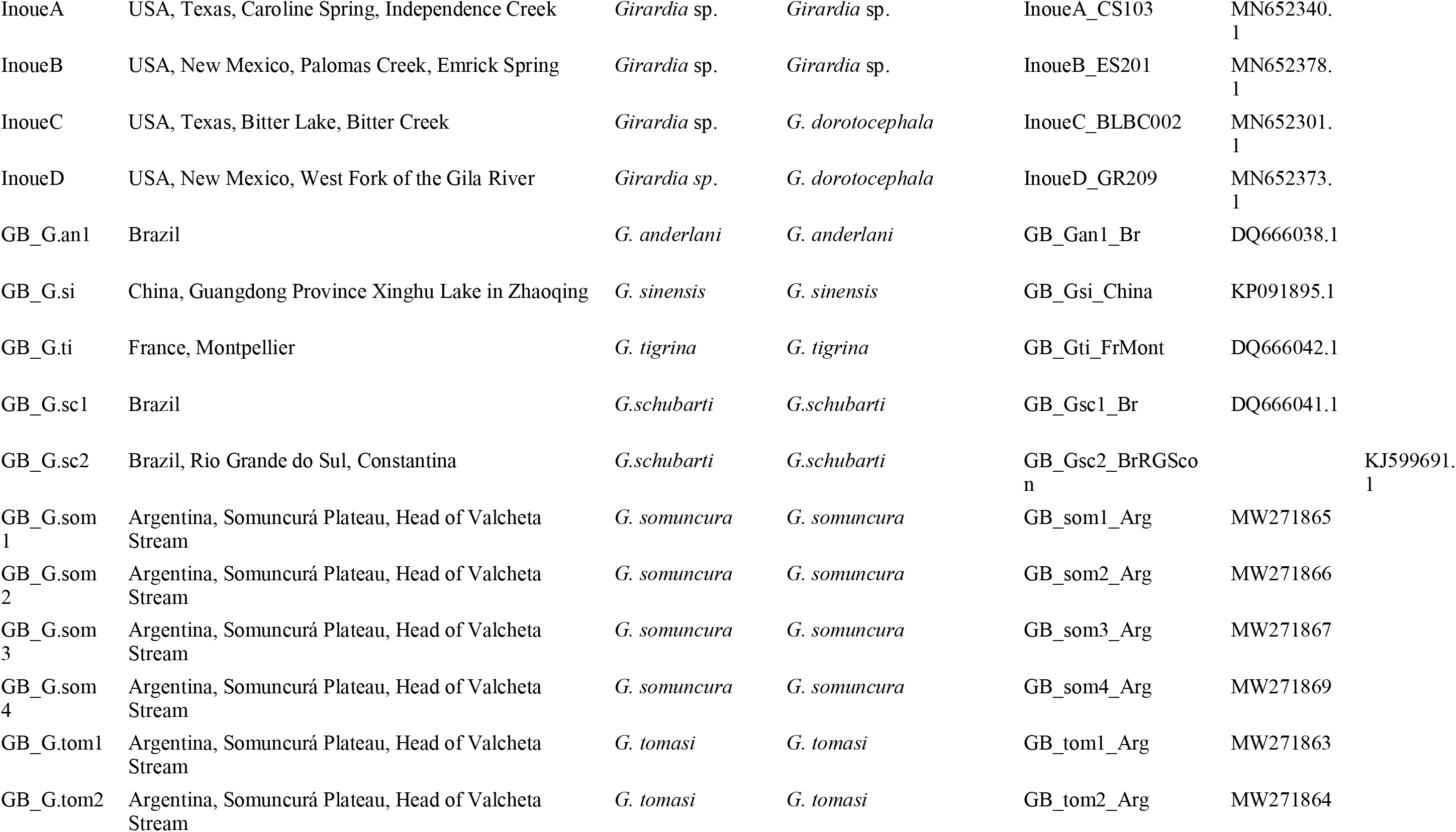
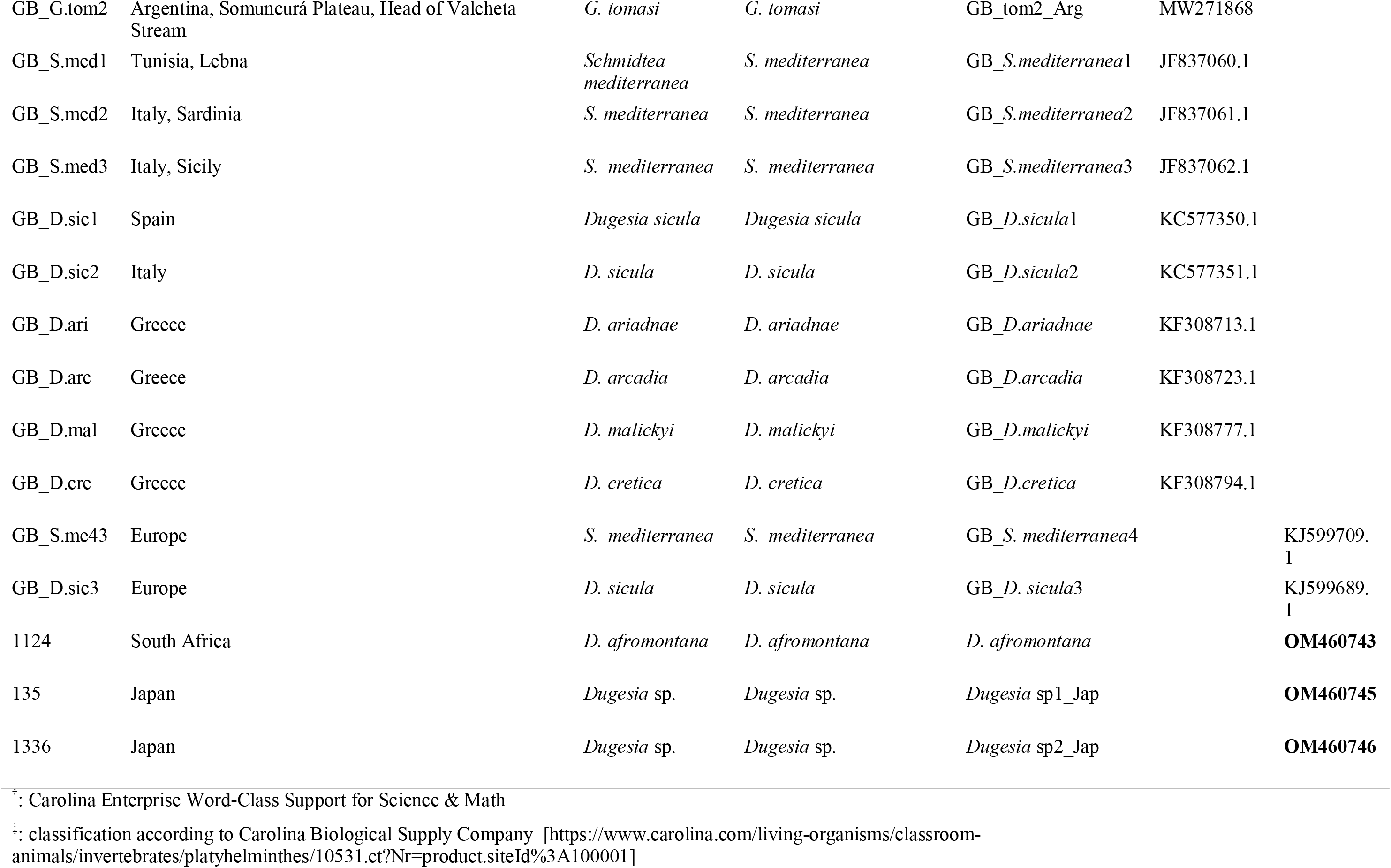

## Appendix 2

***Girardia clandistina* Sluys & Benítez-Álvarez, sp. nov.**

**Holotype:** Naturalis Biodiversity Center, ZMA V.Pl. 976.4, Arroyo Saves Dept., Canalos BS, Uruguay, 1-3 January 1987, sagittal sections on 6 slides.

**Etymology:** The specific epithet is based on the Latin adjective *clandistinus*, secret, concealed, and alludes to the fact that it concerns a sibling species.

**Differential diagnosis:**

A species of *Girardia* with low triangular head with bluntly pointed tip and short, broad auricles. Dorsal body colouration variable, being of a reticulated type with darkish spots and also a pair of dark stripes, separated by a pale mid-dorsal streak, or composed of a dark background interspersed with white splotches and with a pale middorsal line, or variations on these two major patterns. Reproductive complex basically as in *G. tigrina*, the only, but consistent, anatomical difference between the two species residing in the coat of muscles around the bursal canal. In North American *G. tigrina* this coat of muscles is simple, consisting of a thin subepithelial layer of circular muscle, followed by an equally thin layer of longitudinal muscle fibres. In contrast, *G. clandistina* possesses a bursal canal musculature that consists of a well-developed coat of intermingled circular and longitudinal muscle fibres.

