## Supplementary figures S1 and S2 for "First molecular phylogeny of the freshwater planarian genus *Girardia* (Platyelminthes, Tricladida) unveils hidden taxonomic diversity and initiates resolution of its historical biogeography"

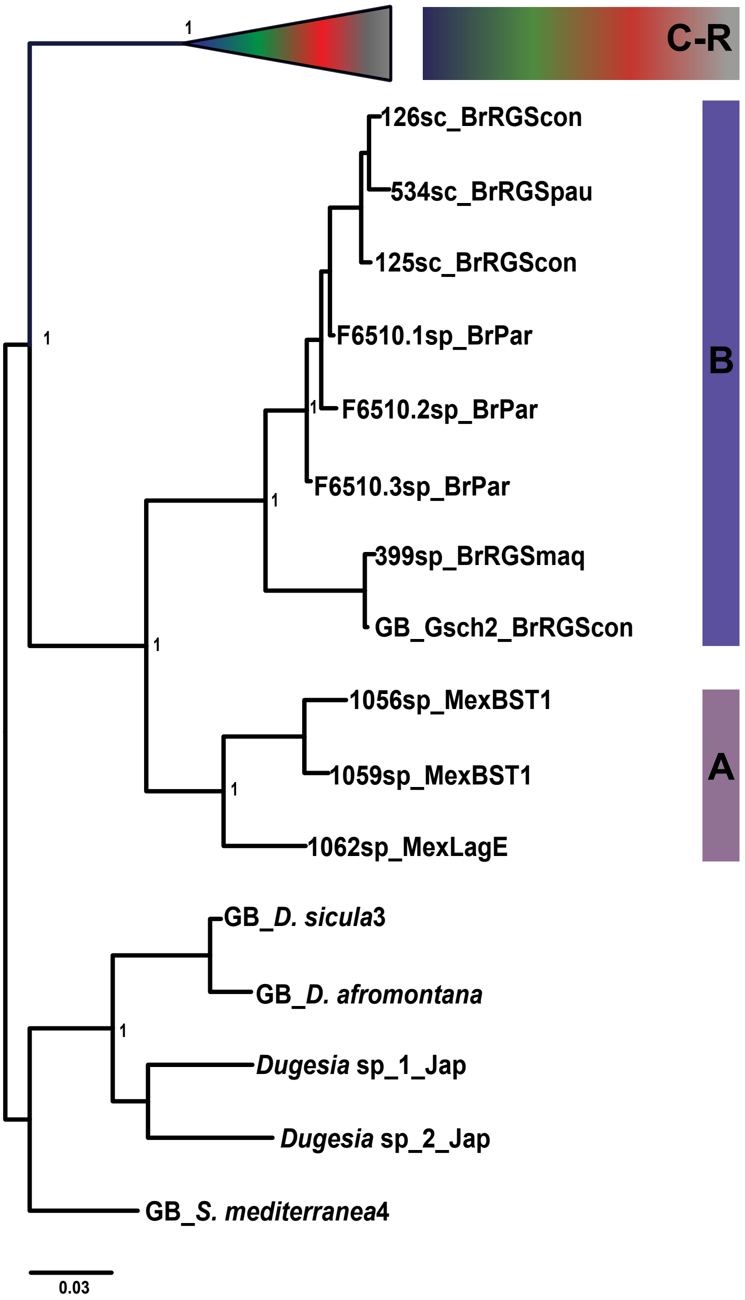


Figure S1. Bayesian Inference tree from Dataset4 (EF1a with outgroup). The sister group of *Girardia schubarti* and unclassified samples from Mexico have been collapsed. Values at nodes correspond to posterior probability support. Scale bar: number of substitutions per nucleotide position.

Figure S2. Bayesian Inference trees from A: Dataset1 (COI withouth outgroup) and B: Dataset3 (EF1a without outgroup). Values at nodes correspond to posterior probability support. Scale bar: number of substitutions per nucleotide position.

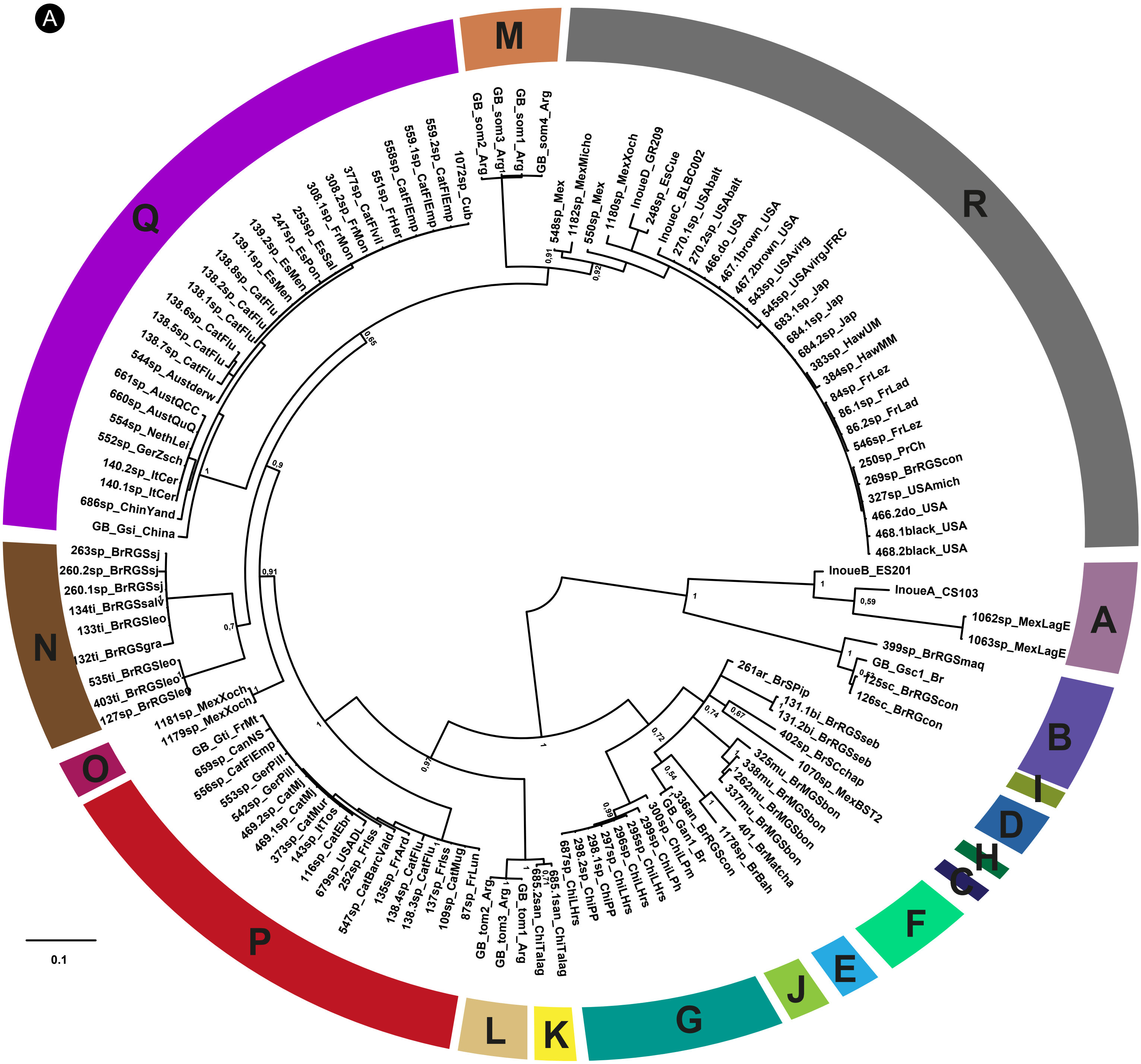


### Phylogenetic clades

- **R:** ***G. dorotocephala*** France, Japan, Hawaii, USA, Mexico, Brazil
- **Q:** ***G. sinensis*** China, Spain, France, Italy, Germany,Netherlands, Australia, Cuba
- **P:** ***G. tigrina*** USA, Canada, France, Spain, Italy, Germany
- **O:** ***Girardia* sp.** Mexico
- **N:** ***G. tigrina*** Brazil
- **M:** ***G. somuncura*** Argentina
- **L:** ***G. tomasi*** Argentina
- **K:** ***G. sanchezi*** Chile
- **J:** ***G. anderlani*** Brazil
- **I:** ***G. arenicola*** Brazil
- **H:** ***Girardia* sp.** Brazil
- **G:** ***Girardia* sp.** Chile
- **F:** ***G. multidiverticulata*** Brazil
- **E:** ***Girardia* sp.** Brazil (cave)
- **D:** ***G. biapertura*** Brazil
- **C:** ***Girardia* sp.** Mexico
- **B:** ***G. schubarti*** Brazil
- **A:** ***Girardia* sp.** Mexico, USA

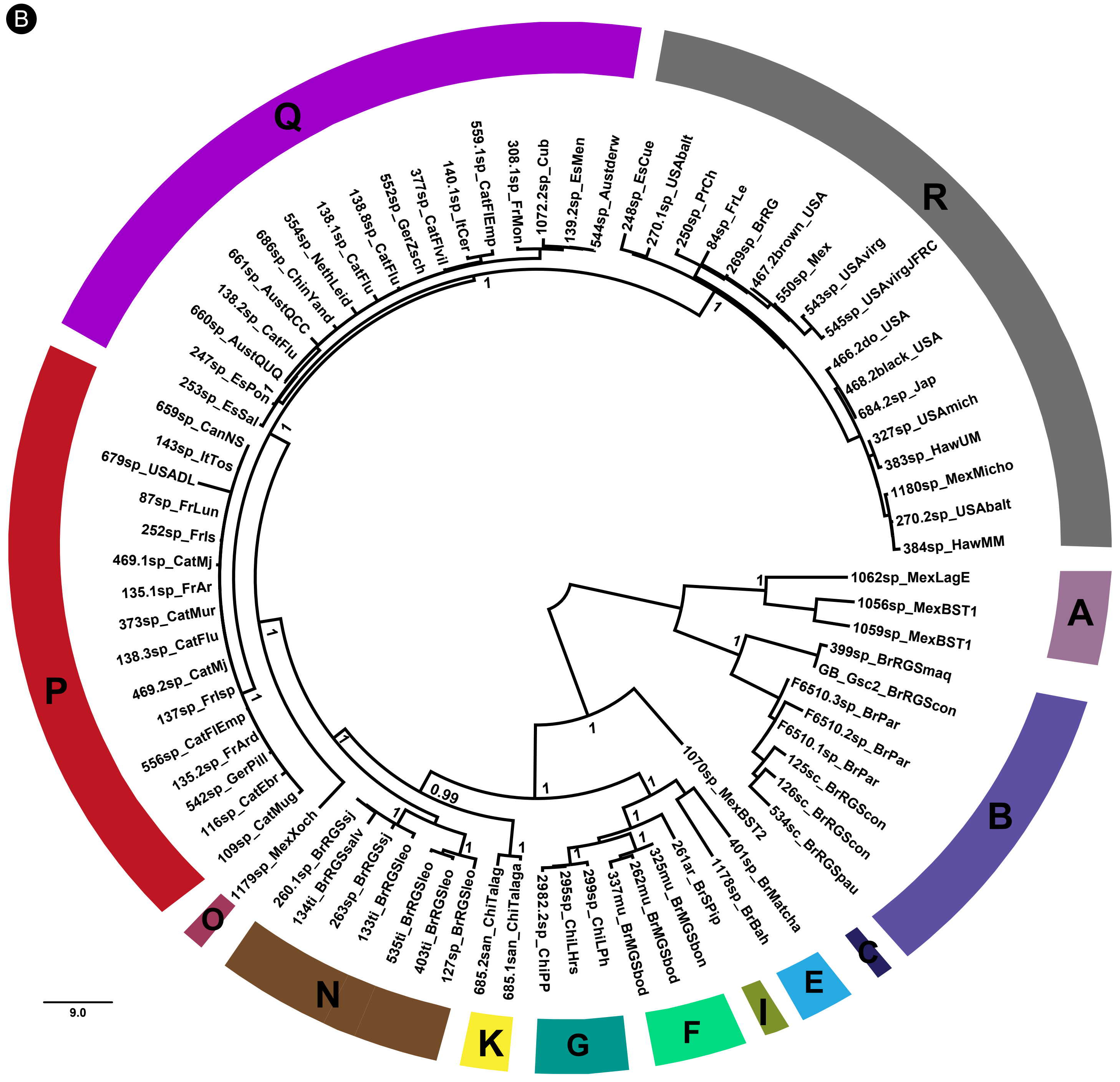


### Phylogenetic clades

- R: ***G. dorotocephala*** France, Japan, Hawaii, USA, Mexico, Brazil
- Q: ***G. sinensis*** China, Spain, France, Italy, Germany, Netherlands, Australia, Cuba
- P: ***G. tigrina*** USA, Canada, France, Spain, Italy, Germany
- O: ***Girardia* sp.** Mexico
- N: ***G. tigrina*** Brazil
- M: ***G. somuncura*** Argentina
- L: ***G. tomasi*** Argentina
- K: ***G. sanchezi*** Chile
- J: ***G. anderlani*** Brazil
- I: ***G. arenicola*** Brazil
- H: ***Girardia* sp.** Brazil
- G: ***Girardia* sp.** Chile
- F: ***G. multidiverticulata*** Brazil
- E: ***Girardia* sp.** Brazil (cave)
- D: ***G. biapertura*** Brazil
- C: ***Girardia* sp.** Mexico
- B: ***G. schubarti*** Brazil
- A: ***Girardia* sp.** Mexico, USA
